# Identification of Cis-Regulatory Sequences Controlling Pollen-Specific Expression of Hydroxyproline-Rich Glycoprotein Genes in *Arabidopsis thaliana*

**DOI:** 10.1101/2020.08.19.256693

**Authors:** Yichao Li, Maxwell Mullin, Yingnan Zhang, Frank Drews, Lonnie Welch, Allan Showalter

## Abstract

Hydroxyproline-rich glycoproteins (HRGPs) are a superfamily of plant cell wall structural proteins that function in various aspects of plant growth and development, including pollen tube growth. We have previously characterized HRGP superfamily into three family members: the hyperglycosylated arabinogalactan-proteins, the moderately glycosylated extensins, and the lightly glycosylated proline-rich proteins. However, the mechanism of pollen-specific HRGP expression remains untouched. To this end, we developed an integrative analysis pipeline combining RNA-seq gene expression and promoter sequences that identified 15 transcriptional cis-regulatory motifs responsible for pollen-specific expression of HRGP in *Arabidopsis Thaliana*. Specifically, we mined the public RNA-seq datasets and identified 13 pollen-specific HRGP genes. Ensemble motif discovery with various filters identified 15 conserved promoter elements between *Thaliana* and *Lyrata*. Known motif analysis revealed pollen related transcription factors of GATA12 and brassinosteroid (BR) signaling pathway regulator BZR1. Lastly, we performed a machine learning regression analysis and demonstrated that the identified 15 motifs well captured the HRGP gene expression in pollen (R=0.61). In conclusion, we performed the integrative analysis as the first-of-its-kind study to identify cis-regulatory motifs in pollen-specific HRGP genes and shed light on its transcriptional regulation in pollen.

## Introduction

Tissue-specific gene expression patterns are maintained by the combinatorial binding of transcription factors (TFs) to DNA motifs in a cooperative and competitive manner. DNA motifs are specific short DNA sequences, often 8-20 nucleotides in length[1], which are statistically overrepresented in a given set of sequences. Extensive studies have been done to characterize regulatory factors and sequences responsible for tissue-specific gene expression in human and mouse[2] [3]. However, to our knowledge, there is no such studies that elucidate transcriptional regulatory motifs responsible for pollen-specific gene expression, particularly for Hydroxyproline-rich glycoproteins (HRGPs) in *Arabidopsis thaliana.*

Hydroxyproline-rich glycoproteins (HRGPs) are a superfamily of plant cell wall proteins involved in various aspects of plant growth and development[4]. The HRGP superfamily consists of three family members, the extensins (EXTs), arabinogalactan-proteins (AGPs), and proline-rich proteins (PRPs). Although all HRGPs contain hydroxyproline, the three family members are distinguished by their unique amino acid compositions, repeated amino acid motifs, and the degree and type of glycosylation. For example, AGPs can be identified by their biased amino acid compositions of Pro (P)/Hyp (O), Ala (A), Ser (S), and Thr (T), their frequent occurrence of AP and PA dipeptide repeats, and their large arabinose and galactose-rich polysaccharide chains attached to O residues. In contrast, EXTs tend to be rich in S, P/O, Val (V), Tyr (Y), and Lys (K), SOOOO pentapeptide repeats, and have multiple short arabinose oligosaccharide side chains attached to their O residues. Finally, PRPs are rich in P, V, K, Cys (C), and T, often have various Pro/Hyp-rich amino acid repeat motifs in which not all the P residues are modified to form O and are the least glycosylated members of the HRGP superfamily.

Bioinformatic programs analyzing genomic/proteomic data from the model genetic plant, *Arabidopsis thaliana*, have identified 166 HRGPs consisting of 85 AGPs, 59 EXTs, 18 PRPs, and 4 AGP/EXT hybrid HRGP[4]. Most HRGP genes are found to be expressed in a tissue-specific manner. Virtually all HRGP genes are significantly differentially expressed in biotic and abiotic stress conditions. This information has provided new insight to the HRGP superfamily and is being used by researchers to facilitate and guide further research in the field. However, one of the untouched questions is the transcriptional regulation of HRGP genes, particularly in mature pollen (e.g., sperm cells).

To fill the void, in this study, we analyzed 113 RNA-seq data from Araport11[5] and identified 13 pollen-specific HRGP genes based on tissue-specificity index (Tau[6]). Ensemble motif discovery was performed using Emotif-Alpha[7] and identified 15 pollen-specific *de novo* motifs. Known motif matching based on PlantTFDB[8] and TOMTOM[9] identified interesting TFs that have been previously reported in pollen, such as GATA12 and BZR1. Regression analysis between HRGP gene expression and the identified motifs showed a reasonable correlation. Our results provide the first overview of the putative cis-regulatory elements in pollen-specific HRGP genes.

## Results

### Integrative analysis pipeline for pollen-specific HRGP and promoter motifs

To answer the question of what cis-regulatory motifs control pollen-specific HRGP gene expression, we developed a systematic bioinformatic pipeline for interactive analysis of pollen-specific HRGP genes and *de novo* promoter motifs (Figure 1) based on several published databases and tools, including gene expression database Araport11[5], known transcription factor binding sites database plantTFDB[8], and motif discovery tool Emotif-Alpha[7], which is an ensemble motif discovery pipeline that integrates more than 10 different motif discovery tools, such as GimmeMotifs[10], DECOD[11], and DME[12]. We obtained 166 HRGP genes from [4], in which 13 genes were defined as pollen-specific and 132 genes were defined as non-pollen (i.e., not expressed in pollen) based on public RNA-seq datasets. We then performed ensemble motif discovery using Emotif-Alpha and defined a relaxed set of motifs (i.e., filter A) and a rigorous set of motifs (i.e., filter B). Lastly, using the results from filter A and filter B, we performed a machine learning regression analysis using the identified 15 motifs (one motif was identified in both filters) and the 166 HRGP genes. The regression model showed that the identified motifs well captured HRGP gene expression in pollen (R=0.61).

**Figure 1.**
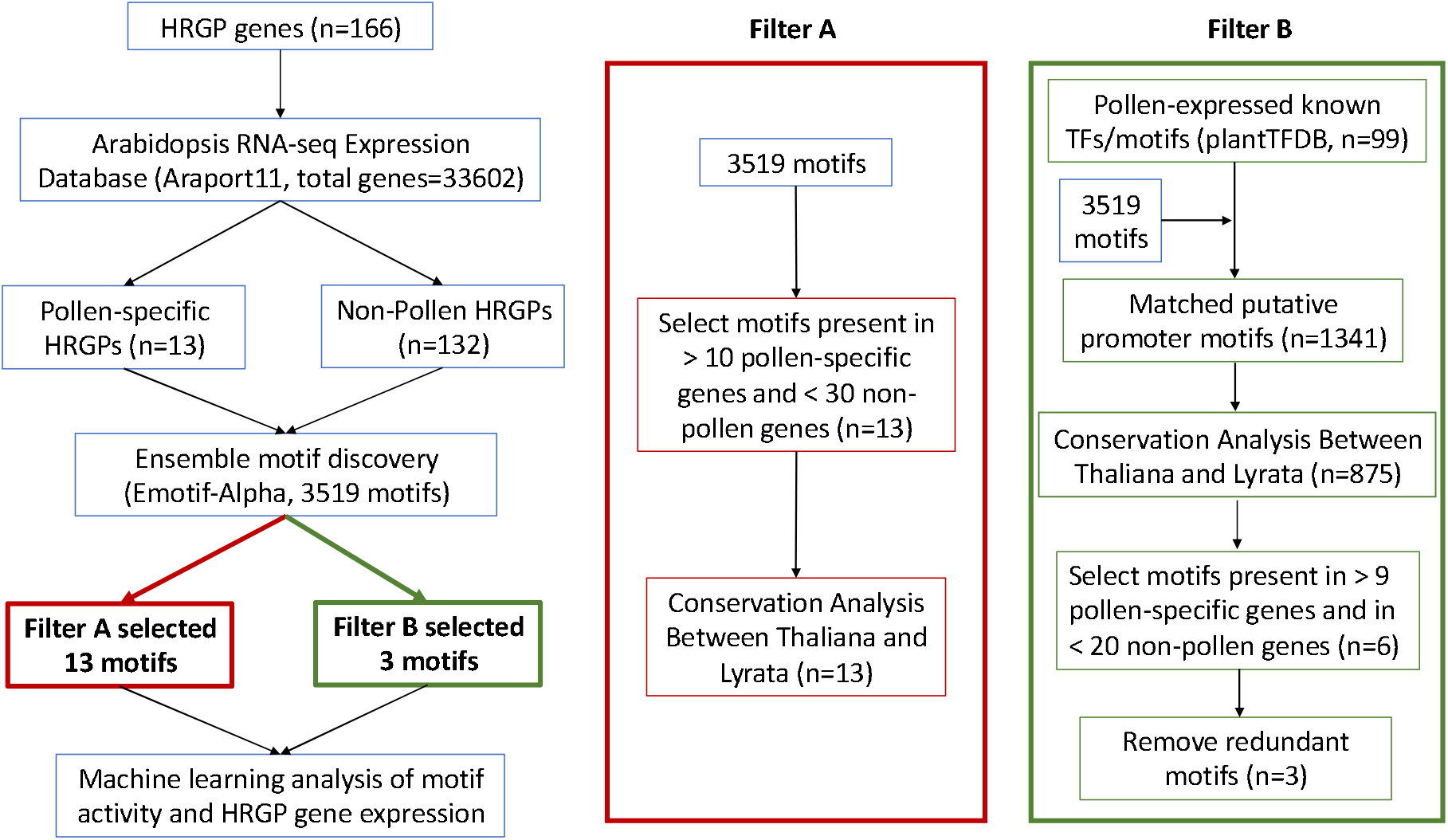
Workflow of the integrative analysis combining HRGP gene expression and promoter sequences. HRGP genes were split into pollen-specific and non-pollen (i.e., not expressed in pollen) based on tissue-specific gene expression analysis from Araport11[5]. Ensemble motif discovery was performed on the aforementioned two sets and totally 3519 motifs were found, which were gone through two filters. Filter A is a more relaxed approach where the motifs were filtered based on the number of occurrences (>10 in pollen-specific HRGPs and <30 in non-pollen HRGPs) and conservation criterion. Filter B is a more rigorous approach: 99 motifs whose cognate transcription factors expressed in pollen were obtained from plantTFDB[8]. Known motif similarity found that 1341 *de novo* motifs out of the total 3519 motifs were highly similar to the 99 motifs (p-value<0.001). The set was then filtered by conservation, number of occurrences, and redundant motifs were removed. In total, filter A and filter B identified 15 motifs controlling pollen-specific HRGP gene expression, which were then fit into a regression model that integrated the promoter elements and HRGP gene expression.

**Figure 2.**
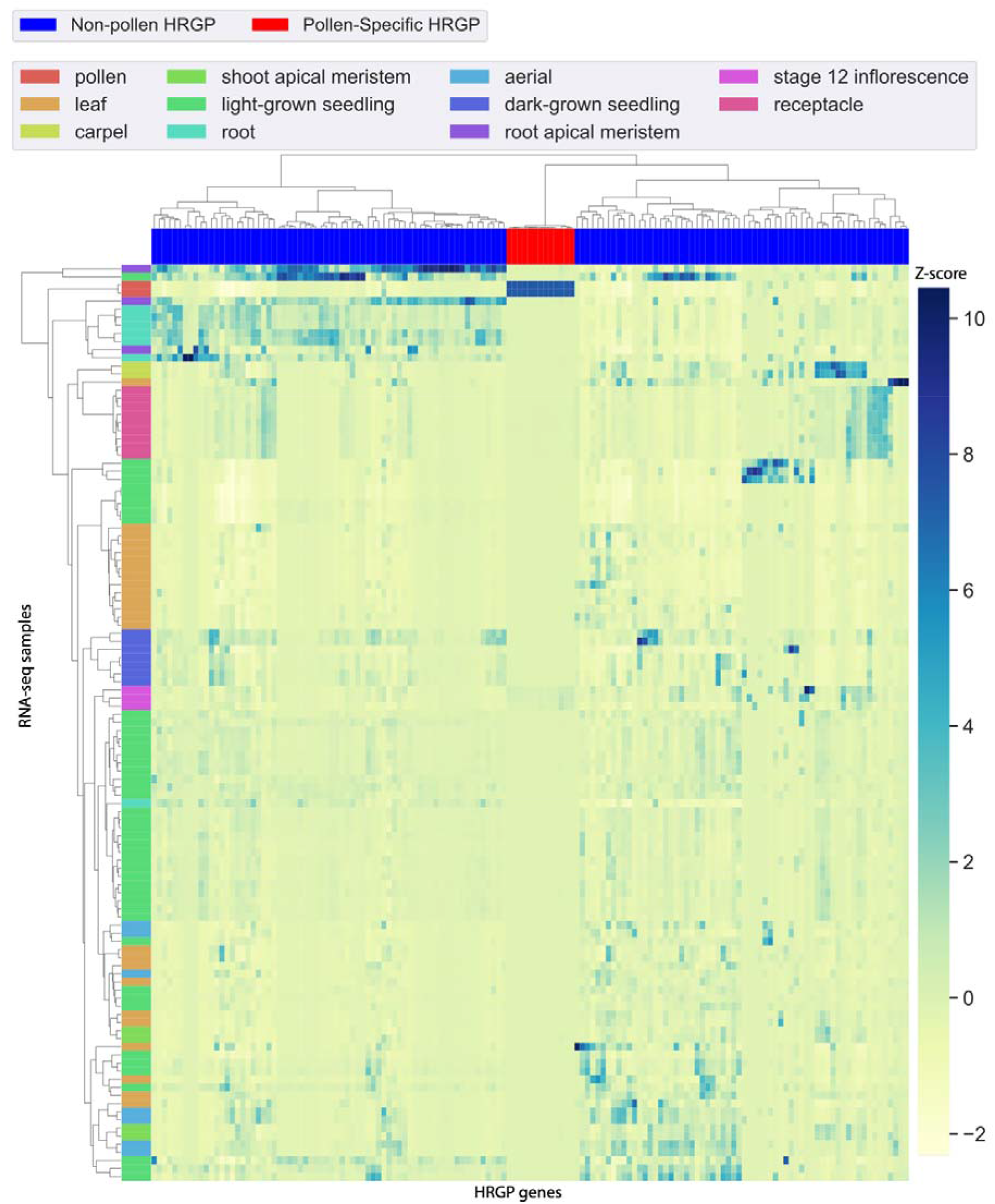
Gene expression heatmap of 13 pollen-specific and 132 non-pollen HRGPs. Gene expression z-scores were calculated by seaborn.clustermap[14] where the larger value means higher expression. Row represents 113 RNA-seq datasets from 11 different tissues. Column represents the HRGP genes.

**Figure 3.**
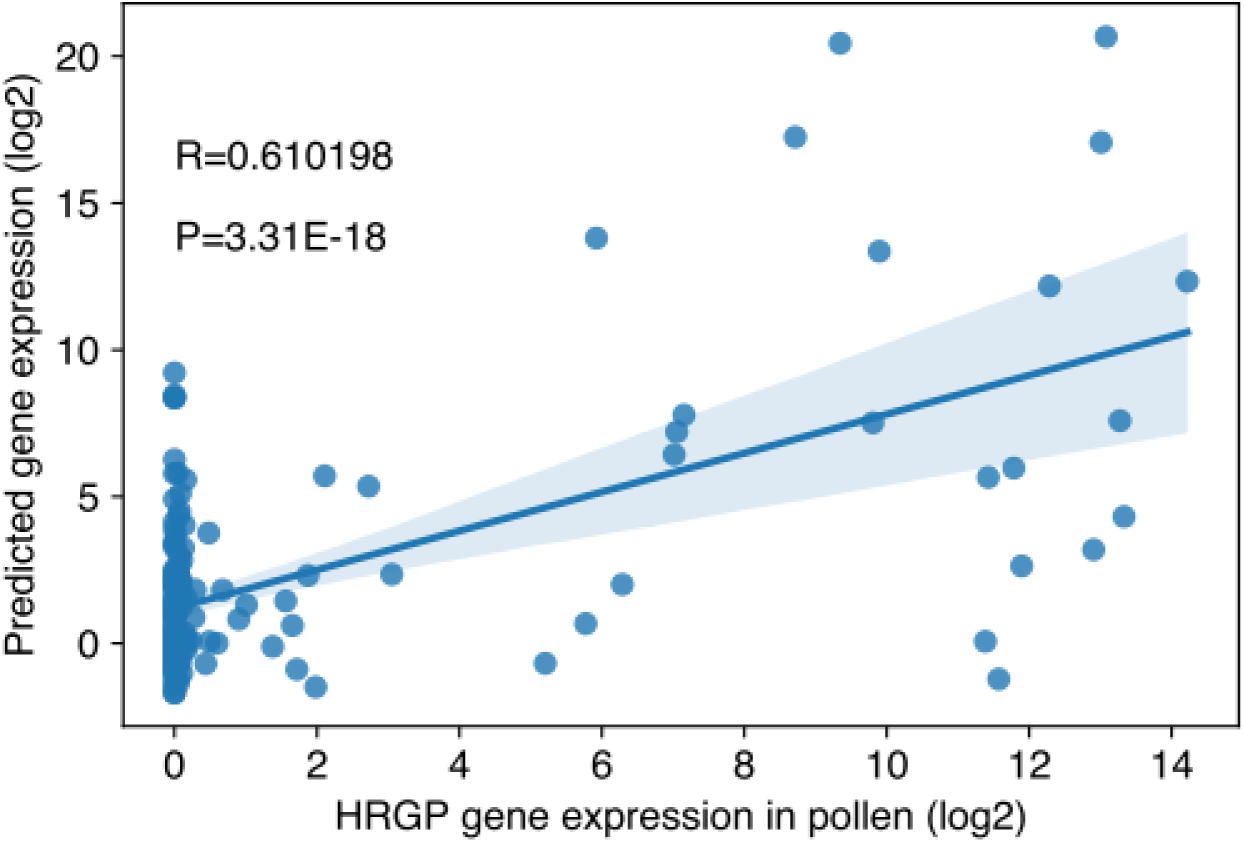
Machine learning regression analysis using the identified 15 pollen-specific motifs. Each point is an HRGP gene. X axis is the mean HRGP gene expression in pollen and the Y axis is the predicted gene expression value. Pearson correlation coefficient is 0.61 and the p-value is 3.31E-18.

### Identification of pollen-specific HRGP genes

Tissue-specific gene expression analysis was performed using 113 RNA-seq samples in 11 different tissues from Araport11[5]. We computed the tissue specificity index (Tau[6]) for each gene. Tau value varies from 0 to 1, where lower Tau means more universally expressed genes and higher Tau means more tissue-specifically expressed genes. A gene is defined as tissue-specific if its Tau value is greater than 0.85. Thus, a HRGP gene is called pollen-specific if Tau > 0.85 and the maximal expressed tissue is pollen (Table S1). Using these criteria, we have identified 13 pollen specific HRGPs, including 8 EXTs and 5 AGPs. To identify promoter motifs controlling pollen-specific HRGP gene expression, we further defined a background set of 132 non-pollen HRGP genes for discriminative motif discovery using Emotif-Alpha[7]. Gene expression heatmap showed that the pollen-specific HRGPs were almost exclusively expressed in pollen and non-pollen HRGPs were expressed in other tissues but not in pollen (Figure 1). We notice that 12 pollen-specific HRGPs have been previously reported to be pollen-specific by an analysis of gene expression microarrays in [4]. AT1G54215 (EXT32) is a newly identified pollen-specific HRGP using public RNA-seq datasets. Due to cross-hybridization and saturation of signals, microarrays have limited detection capability and high background noises. RNA-seq, on the other hand, is far more sensitive and precise[13]. Thus, we have identified one additional pollen-specific HRGP comparing to [4].

### Motif discovery filter A: a relaxed set of pollen-specific HRGP motifs

Next, we performed ensemble motif discovery using Emotif-Alpha[7] on the 13 pollen-specific HRGP genes against the 132 non-pollen HRGP genes. Gene promoters (1kb upstream of the translation start site) were retrieved from Ensembl Biomart[15]. Emotif-Alpha integrated 11 motif discovery tools and we have found 3519 motifs in total.

Filter A selected motifs presented in 11 (85%) or more pollen specific HRGP genes and presented in at most 30 (23%) non-pollen HRGP genes. This returned 13 motifs, which were also conserved in *Arabidopsis Lyrata.* Table 1 showed the identified 13 motifs. Interestingly, we found that 4 out of the 13 motifs matched to known motifs based on TOMTOM[9] (p-value < 0.001).

**Table 1.**
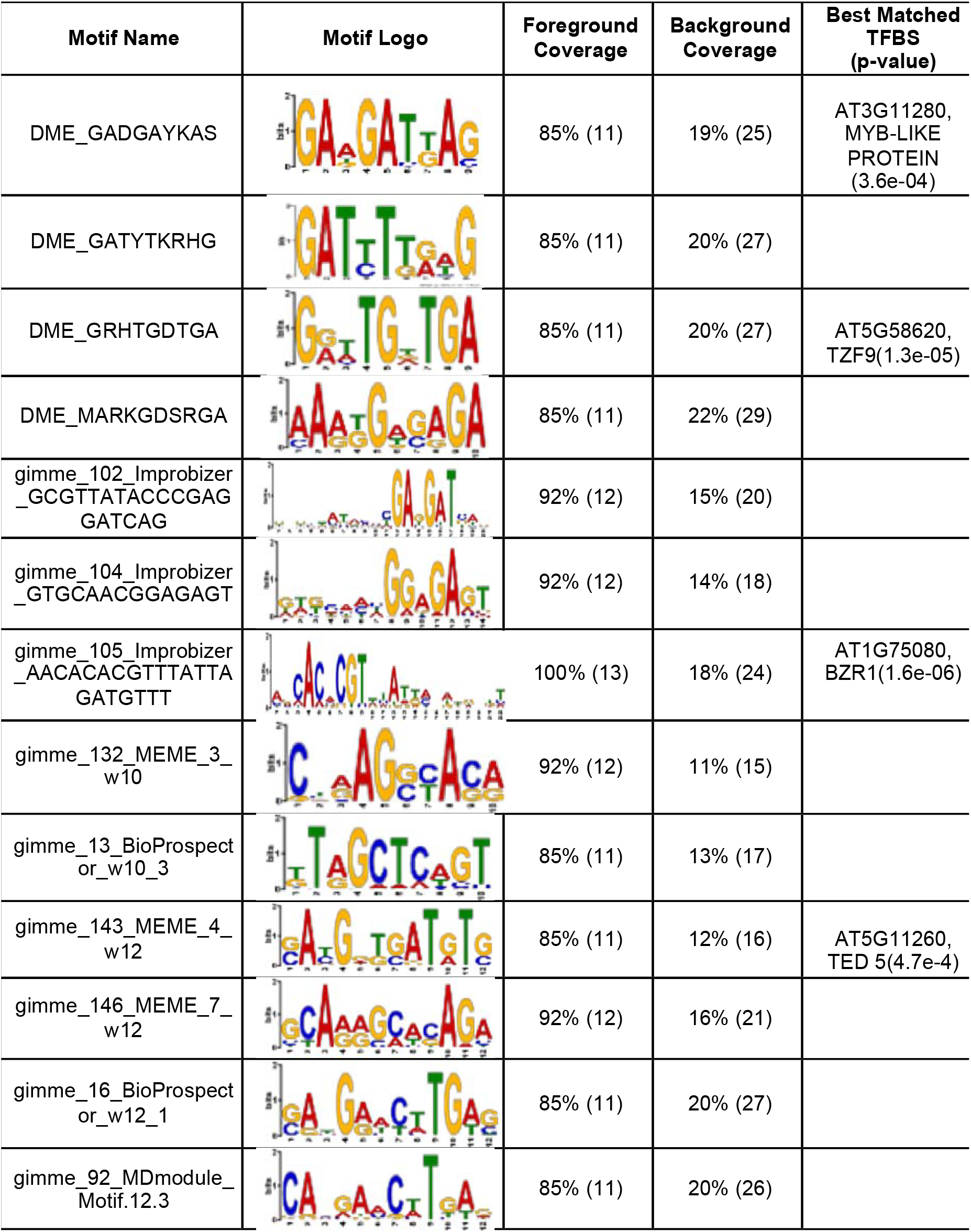
Motifs identified in Filter A. The foreground coverage indicates the percentage (and number) of pollen-specific HRGP genes having that motif in its promoter region. The background coverage is the percentage (and number) of the non-pollen HRGP genes having that motif in its promoter region. The relative frequency is the foreground coverage divided by the background coverage. The matched TFBS p-value is a known transcription factor binding site that relates to pollen along with the accompanying p-value.

**Table 2.**
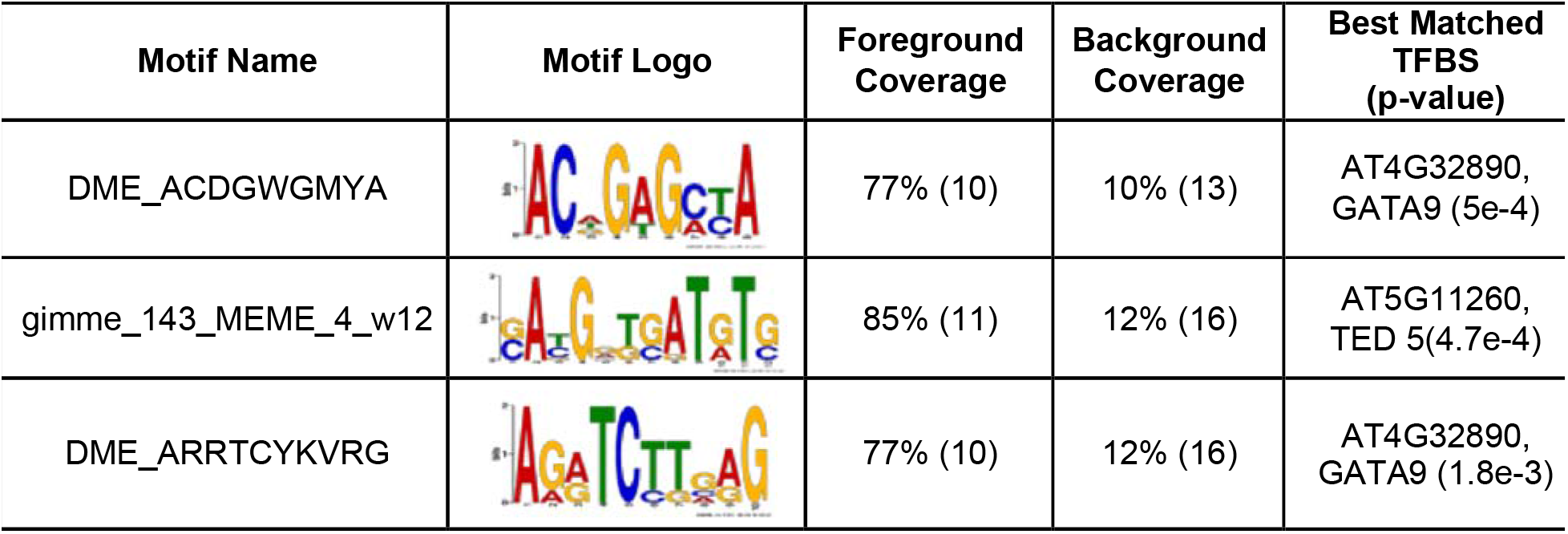
Motifs identified in Filter B. The foreground coverage indicates the percentage (and number) of pollen-specific HRGP genes having that motif in its promoter region. The background coverage is the percentage (and number) of the non-pollen HRGP genes having that motif in its promoter region. The relative frequency is the foreground coverage divided by the background coverage. The matched TFBS p-value is a known transcription factor binding site that relates to pollen along with the accompanying p-value.

It is worth noting that motif gimme_105_Improbizer AACACACGTTTATTAGATGTTT occurred in all 13 pollen-specific HRGP genes and this motif is highly similar (p-value = 1.6e-06) to the known BZR1 (Brassinazole Resistant 1) binding motif. Brassinosteroid (BR) is an important class of steroid hormones in plants that regulates gene expression and cell development[16]–[18]. BZR1 is a key transcription factor in the BR signaling pathway, where the binding of BR to a cell surface receptor kinase (BRI1) directly regulates the phosphorylation of BZR1, which then binds to the promoters of BR responsive genes. BR is first discovered in pollen where it regulates cell elongation. Although it is later found in all tissues, the highest abundance is found primarily in pollen, seeds and fruit[16]. The discovery of BZR1-like binding site in the promoters of pollen-specific HRGP suggests that these HRGPs are likely to be regulated by the BR signaling pathway. Indeed, cell wall modification is reported to be one of the major functions targeted by the BR pathway[16].

### Motif discovery filter B: a rigorous set of pollen-specific HRGP motifs

Filter B is a more rigorous approach where pollen expressed known TFs (i.e., log2 gene expression >=1) and their binding motifs were used. First, 99 motifs whose cognate transcription factors expressed (based on Araport11) in pollen were obtained from plantTFDB[8]. Known motif similarity found that 1341 *de novo* motifs out of the total 3519 motifs were highly similar to the 99 motifs (TOMTOM[9], P<0.001). The set was then filtered by conservation, number of occurrences, and redundant motifs were removed. Motif gimme_143_MEME_4_w12 was also found in filter A approach.

Interestingly, two motifs (i.e., DME_ACDGWGMYA and DME_ARRTCYKVRG) matched to GATA9 binding sites. Since GATA9 is the closest homolog of GATA12 and it is known that “levels of GATA12 were high in mature pollen grains but diminished in the germinated pollen grains and their pollen tubes” [19], it suggests that these two motifs might play a gene regulation role in pollen-specific expression in *Arabidopsis thaliana.*

### Modeling HRGP gene expression in pollen

To investigate how well our identified pollen-specific motifs can predict HRGP gene expression in pollen, we conducted a regression analysis using a well-known machine learning package scikit-learn[20] in Python. The 15 identified motifs were mapped to the 166 HRGP gene promoters and FIMO[21] motif mapping p-value (log10 transformed) was used as features. A gradient booting tree was trained and evaluated using 3-fold cross validation[20]. We obtained a good correlation between true HRGP gene expression and predicted values (R=0.61, P=3.31E-18), suggesting that the 15 identified motifs might control HRGP gene expression in pollen.

## Discussion

Tissue-specific gene expression analysis coupled with motif discovery is the first step to understand the regulatory mechanism of tissue specificity. In this study, we applied the tissue-specificity index and defined 13 pollen-specific HRGP genes. Using two different filters, including both relaxed and rigorous criteria, we identified 15 pollen-specific motifs. Regression analysis showed that the identified motifs can be used to predict HRGP gene expression in pollen. However, other factors, such as chromatin openness, histone modifications, and DNA methylation are missing in our current model due to limited public data resources in pollen tissue in *Arabidopsis Thaliana.* To overcome this limit, ATAC-seq and ChlP-seq can be performed to further explore the transcriptional regulatory landscape of HRGP genes as well as other genes in pollen.

## Materials and Methods

### Characterization of pollen-specific HRGP genes

A list of 33602 Arabidopsis thaliana genes of TAIR ID was downloaded from TAIR website (https://www.arabidopsis.org/download_files/Genes/TAIR10_genome_release/TAIR10_gene_lists/TAIR1_0_gene_type). Then, for each gene in the list, its gene expression profile in 113 RNA-seq experiments was retrieved from Araport11 using the python API, intermine.webservice. The description of the RNA-seq dataset can be found at: https://www.araport.org/rna-seq-read-datasets-used-araport11. The entire gene expression profile can be accessed in Table S1. To determine pollen-specific expression, the tissue specificity index, Tau, was used. In a recent benchmarking comparison, Tau was found to be the most robust and biological relevant method[20]. Tau varies from 0 to 1, where lower Tau means more universally expressed and higher Tau means more tissue specifically expressed. As recommended in [5], genes with Tau > 0.85 were considered tissue-specific. The tissue type was determined by the largest expressed tissue. All expression values were log-transformed before calculating Tau; values <1 were set to 0 after log transformation[20]. Using this method, we have characterized tissue-specific expression patterns for 26500 genes. Data can be accessed in Table S2.

A list of 166 HRGPs was reported by Showalter et al. [3]. Pollen-specific HRGPs are defined as a list of HRGPs that are pollen-specific expressed. Non-pollen HRGPs are defined as a list of HRGPs that have no expressions (expression value is 0 after log transformation) in pollen (Table S3).

### Promoter retrieval and ensemble motif discovery

Promoters of pollen-specific HRGPs and non-pollen HRGPs were retrieved from Ensembl Plant v35 Biomart web interface using gene stable ID, Flank (Gene) Coding Region, and Upstream flank 1000 bp.

To identify regulatory motifs for pollen-specific HRGPs, Emotif-Alpha, an ensemble motif discovery pipeline was applied. Foreground promoter set was the list of 13 pollen-specific HRGPs. Background promoter set was the list of 132 non-pollen HRGPs. Emotif-alpha has integrated 11 motif discovery tools: GimmeMotifs, MEME, Weeder, BioProspector, AMD, Homer, GADEM, MDmodule, and Improbizer, DME, and DECOD. Motif length was set to be 6-16nt. Fimo [21] was used for motif scanning. The discriminative power of the motifs was assessed by a random forest classifier using scikit-learn. Motif similarity was assessed by TOMTOM[9]. Two motifs were similar if their TOMTOM[9] p-value was less than 0.001 and the motif with less occurrences in pollen-specific HRGPs would be filtered out.

### Conservation analysis

Conservation analysis was performed using the method adopted by Roy et, al[22]. Orthologous information between *Thaliana* and *Lyrata* were retrieved from Ensembl Plant Biomart v39[15]. CLUSTALW2[23] was used to do multiple sequence alignment with gap open penalty of 10 and extension penalty of 0.1. A motif was defined as conserved if it occurred at the same position in the orthologous promoter alignment.

### Machine learning

The 15 identified motifs were scanned on the 166 HRGP promoters using FIMO[21] and the −log10 motif scanning p-value was used as machine learning features for regression. The regression algorithm was implemented using scikit-learn GradientBoostingRegressor with parameters of subsample=0.3, criterion=“mae”, min_samples_split=5, max_depth=1[20]. The evaluation was performed using 3-fold cross-validation using the KFold function.

## Acknowledgements

We would like to thank Prashant Kuntala and Isaac Torres for their computer programming support. Additionally, we would like to thank Richard Wolfe, Rami Ouran, and Xiao Liu for helpful discussions.

## Author contributions

Y.L, M.M performed the analyses and wrote the article; Y.Z provided biological insights and wrote the article; F.D, A.S, L.W supervised and complemented the writing.

## Data Availability

All data and source code used in this study is available at: https://github.com/YichaoOU/Pollen_specific_motifs

## Supplementary Files

**Table S1.**
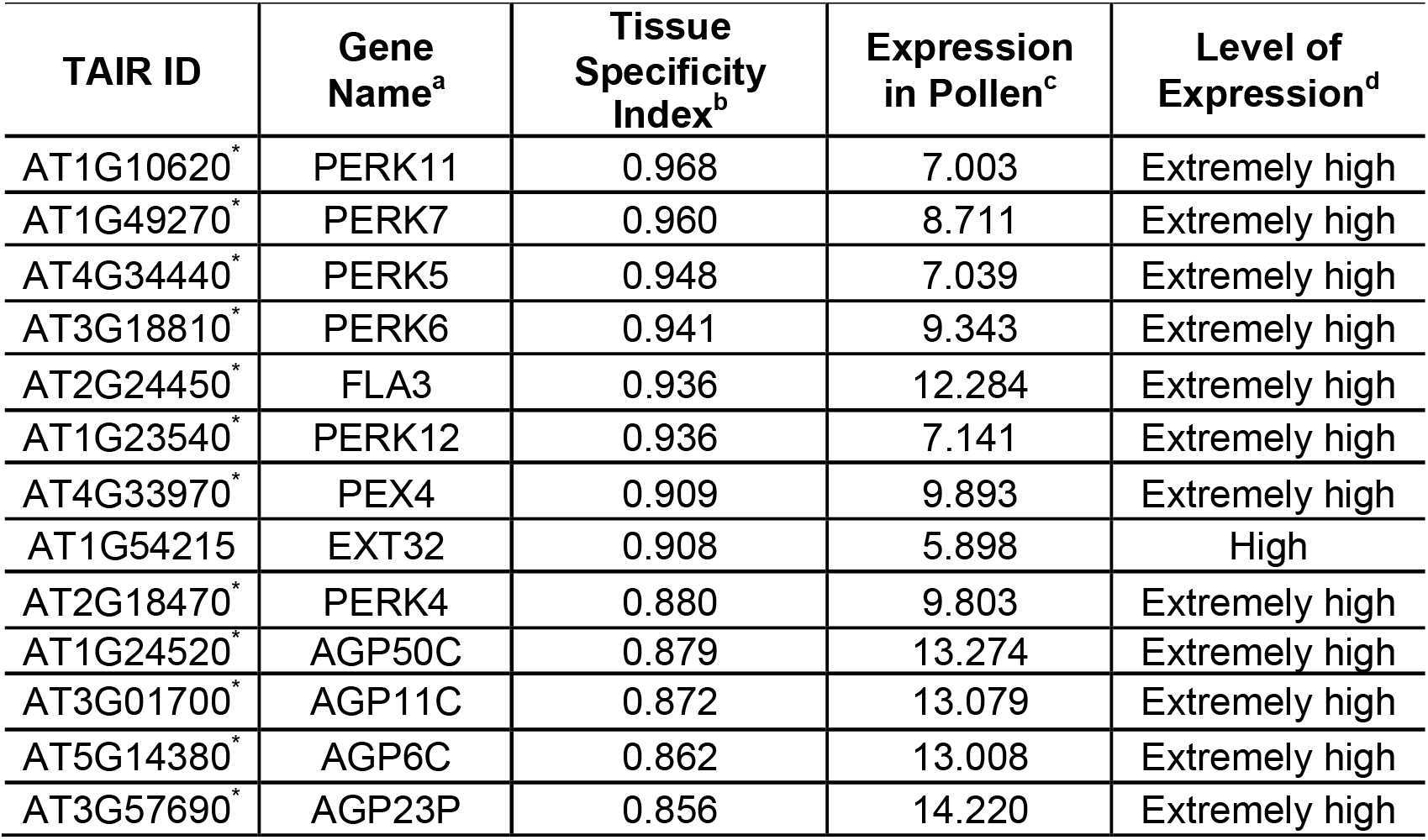
List of pollen-specific HRGP genes. *These genes have been reported to be pollen-specific in [17]. ^a^Gene Name is adopted from [17], where some genes are renamed by the authors to indicate their protein sequence properties. ^b^Tissue specificity index Tau is calculated using the formula presented in [18]. Expression is represented using the median value after log 2 transformation. Expression value is compared to number of standard deviation (std) away from the mean value in all genes’ expression profile in pollen. Extremely high expressed genes have more than 3*std away from the mean and high expressed genes have more than 2*std but less than 3*std away from the mean.

## Reference

[1] F. Zambelli, G. Pesole, and G. Pavesi, “Motif discovery and transcription factor binding sites before and after the next-generation sequencing era,” Brief. Bioinform., vol. 14, no. 2, pp. 225–237, 2012, doi: 10.1093/bib/bbs016.

[2] A. Arvey, P. Agius, W. S. Noble, and C. Leslie, “Sequence and chromatin determinants of celltype–specific transcription factor binding,” Genome Res., vol. 22, no. 9, pp. 1723–1734, 2012, doi: 10.1101/gr.127712.111.

[3] A. Natarajan, G. G. Yardimci, N. C. Sheffield, G. E. Crawford, and U. Ohler, “Predicting cell-type-specific gene expression from regions of open chromatin.,” Genome Res., vol. 22, no. 9, pp. 1711–1722, Sep. 2012, doi: 10.1101/gr.135129.111.

[4] A. M. Showalter, B. Keppler, J. Lichtenberg, D. Gu, and L. R. Welch, “A bioinformatics approach to the identification, classification, and analysis of hydroxyproline-rich glycoproteins.,” Plant Physiol., vol. 153, no. 2, pp. 485–513, Jun. 2010, doi: 10.1104/pp.110.156554.

[5] C.-Y. Cheng, V. Krishnakumar, A. P. Chan, F. Thibaud-Nissen, S. Schobel, and C. D. Town, “Araport11: a complete reannotation of the Arabidopsis thaliana reference genome,” Plant J., vol. 89, no. 4, pp. 789–804, 2017, doi: 10.1111/tpj.13415.

[6] I. Yanai et al., “Genome-wide midrange transcription profiles reveal expression level relationships in human tissue specification,” Bioinformatics, vol. 21, no. 5, p. 650, 2005, doi: 10.1093/bioinformatics/bti042.

[7] A. Grote et al., “Prediction pipeline for discovery of regulatory motifs associated with Brugia malayi molting,” PLoS Negl. Trop. Dis., vol. 14, no. 6, pp. 1–16, 2020, doi: 10.1371/journal.pntd.0008275.

[8] J. Jin et al., “PlantTFDB 4.0: toward a central hub for transcription factors and regulatory interactions in plants,” Nucleic Acids Res., vol. 45, no. D1, p. D1040, 2017, doi: 10.1093/nar/gkw982.

[9] S. Gupta, J. A. Stamatoyannopoulos, T. L. Bailey, and W. S. Noble, “Quantifying similarity between motifs,” Genome Biol., vol. 8, no. 2, p. R24, 2007, doi: 10.1186/gb-2007-8-2-r24.

[10] S. J. van Heeringen and G. J. C. Veenstra, “GimmeMotifs: a de novo motif prediction pipeline for ChIP-sequencing experiments,” Bioinformatics, vol. 27, no. 2, pp. 270–271, 2010, doi: 10.1093/bioinformatics/btq636.

[11] P. Huggins et al., “DECOD: fast and accurate discriminative DNA motif finding,” Bioinformatics, vol. 27, no. 17, pp. 2361–2367, 2011, doi: 10.1093/bioinformatics/btr412.

[12] A. D. Smith, P. Sumazin, and M. Q. Zhang, “Identifying tissue-selective transcription factor binding sites in vertebrate promoters,” Proc. Natl. Acad. Sci., vol. 102, no. 5, pp. 1560–1565, 2005, doi: 10.1073/pnas.0406123102.

[13] Z. Wang, M. Gerstein, and M. Snyder, “RNA-Seq: a revolutionary tool for transcriptomics,” Nat Rev Genet, vol. 10, no. 1, pp. 57–63, Jan. 2009.

[14] M. Waskom et al., “mwaskom/seaborn: v0.8.1 (September 2017).” Zenodo, Sep. 2017, doi: 10.5281/zenodo.883859.

[15] R. J. Kinsella et al., “Ensembl BioMarts: a hub for data retrieval across taxonomic space.,” Database (Oxford)., vol. 2011, p. bar030, 2011, doi: 10.1093/database/bar030.

[16] J.-Y. Zhu, J. Sae-Seaw, and Z.-Y. Wang, “Brassinosteroid signalling,” Development (Cambridge, England), vol. 140, no. 8. pp. 1615–1620, Apr. 2013, doi: 10.1242/dev.060590.

[17] Q. Ye et al., “Brassinosteroids control male fertility by regulating the expression of key genes involved in Arabidopsis anther and pollen development,” Proc. Natl. Acad. Sci., vol. 107, no. 13, pp. 6100–6105, 2010, doi: 10.1073/pnas.0912333107.

[18] Z.-Y. Wang et al., “The brassinosteroid signal transduction pathway,” Cell Res, vol. 16, no. 5, pp. 427–434.

[19] P. Ravindran, V. Verma, P. Stamm, and P. P. Kumar, “A Novel RGL2-DOF6 Complex Contributes to Primary Seed Dormancy in Arabidopsis thaliana by Regulating a GATA Transcription Factor.,” Mol. Plant, vol. 10, no. 10, pp. 1307–1320, Oct. 2017, doi: 10.1016/j.molp.2017.09.004.

[20] F. Pedregosa et al., “Scikit-learn: Machine Learning in {P}ython,” J. Mach. Learn. Res., vol. 12, pp. 2825–2830, 2011.

[21] C. E. Grant, T. L. Bailey, and W. S. Noble, “FIMO: scanning for occurrences of a given motif,” Bioinformatics, vol. 27, no. 7, pp. 1017–1018, Apr. 2011, doi: 10.1093/bioinformatics/btr064.

[22] S. Roy, M. Kagda, and H. S. Judelson, “Genome-wide Prediction and Functional Validation of Promoter Motifs Regulating Gene Expression in Spore and Infection Stages of Phytophthora infestans,” PLOS Pathog., vol. 9, no. 3, pp. 1–18, 2013, doi: 10.1371/journal.ppat.1003182.

[23] M. A. Larkin et al., “Clustal W and Clustal X version 2.0,” Bioinformatics, vol. 23, no. 21, pp. 2947–2948, 2007, doi: 10.1093/bioinformatics/btm404.

